# *Chlamydomonas reinhardtii* exhibits stress memory in the accumulation of triacylglycerols induced by nitrogen deprivation

**DOI:** 10.1101/2021.07.23.453471

**Authors:** Pawel Mikulski, Javier Santos-Aberturas

## Abstract

Stress memory is a phenomenon whereby exposure to initial stress event influences a response to subsequent stress exposures. Studying stress memory is important to understand the cellular behaviour in dynamic environment, especially nowadays, in times with growing environmental instability. Stress memory has been characterized in vascular plants but its occurrence in non-vascular plant species has been rarely investigated.

We hypothesized that stress memory occurs in non-vascular plants in relation to metabolic stress. We sought to test it using accumulation of lipids (triacylglycerols) in model green alga *Chlamydomonas reinhardtii* subjected to nitrogen deprivation stress as a model system.

Here, we established stress memory protocol on *C. reinhardtii* cells. Using a blend of microscopy and gas chromatography methods, we showed that the cells exposed to recurrent stress show differential accumulation of triacylglycerols on the quantitative level without qualitative changes in lipid composition, comparing to single stress controls.

Overall, our results suggest that metabolic stress memory does occur in non-vascular plant *C. reinhardtii* and provides a starting point to characterize metabolic stress memory mechanism. Due to the commercial potential of algae, our findings are relevant for basic science, as well as industrial production of algae-derived compounds.

## Introduction

Stress memory is a phenomenon whereby exposure to initial stress event influences a response to subsequent stress exposures. Its importance is highlighted especially nowadays, in times with growing environmental instability (IPCC, 2018). Stress memory has been characterized in vascular plants (Mozgova *et al.*, 2019), where it is related to, i.e., response to drought (Ding *et al.*, 2012) or temperature (Brzezinka *et al.*, 2016). However, the occurrence of stress memory has been only rarely reported in non-vascular plants (Widiez *et al.*, 2014; Korkaric *et al.*, 2015), and still remains a largely unexplored niche.

Unicellular algae form a powerful biotechnological warehouse for production of chemical compounds. Triacylglycerols (TAGs) are lipids that can be used as the precursors in biodiesel production for academic and industrial research. Model green alga, *Chlamydomonas reinhardtii*, accumulates TAGs in stress conditions such as nitrogen deprivation (Park *et al.*, 2015). TAGs accumulation is reversible (Roustan *et al.*, 2017) and chromatin-based (Ngan *et al.*, 2015), features that make this process a potential subject for stress priming.

How TAGs’ accumulation behaves in *C. reinhardtii* subjected to multiple nitrogen deprivation events and whether the species exhibits metabolic stress priming has not been investigated. We hypothesized that stress memory occurs in the accumulation of TAGs in *C. reinhardtii* subjected to nitrogen deprivation. Here, we established a stress priming growth protocol in *C. reinhardtii*. Using a blend of microscopy and gas chromatography, we showed that algal cells exposed to recurrent stress show differential accumulation of triacylglycerols on the quantitative level without qualitative changes in lipid composition, comparing to single stress controls.

This work provides a proof-of-principle evidence that metabolic stress memory does occur in non-vascular plant *C. reinhardtii* and opens a new avenue to understand the mechanisms behind stress memory and cellular response to the fluctuating environment. Due to commercial potential of algae and usage of stress-induced processes for production of biologics, this work offers a contribution to academic and industrial researchers.

## Material and methods

### Growth conditions and growth medium

Wildtype *C. reinhardtii* strain 137c was grown in liquid cultures on TAP medium with agitation at 180 rpm as described previously (Ngan *et al.*, 2015). For nitrogen depletion, TAP medium without nitrogen source was used as described in (Ngan *et al.*, 2015). The cells were washed twice with nitrogen-depleted medium before media change during stress treatment. Alga cultures were kept in growth cabinets (Sanyo-MLR-352-PE) set for 25°C and supplied with LED fluorescent lights (Newlec, NL/18/LED/T8/4/865 & 840).

### Fixation & Nile Red staining

We performed TAGs’ staining with Nile Red dye on fixed cells. We used 3% (v/v) formaldehyde (FA) on 1 ml culture aliquots, incubated for 20 min on ice, spun at 3000 rpm for 5 min and washed twice with phosphate saline buffer (PBS). Final cell pellet was resuspended in 20-50 μL PBS based on estimation from OD600. 6-7 μL of final solution was spread onto microscopy slides, air dried and refrigerated until staining. Staining was done with 5 μg/mL in 0.1 % acetone for 15 min in dark.

### Microscopy and image analysis

Image acquisition was performed in two channels: Ex488/Em520-600 and Ex633/Em635-700 (Ex = excitation, Em = emission), where former corresponds to specific Nile Red signal and the latter to cell autofluorescence. For intensity quantifications, Ex488/Em520-600 channel was used to avoid a crosstalk with autofluorescence channel. For Z-stacks, we used a step size (z-distance between slices) of 0.55-0.59 μm and line averaging at 2. Following parameters for image acquisition were used: scan speed at 400 Hz, pinhole size at 1 Airy unit (AU) and laser power at 16%. Image analysis was performed in Fiji/ImageJ. Nile Red intensity quantification was done in 3 biological replicates per harvesting point and sample type. We quantified intensity from 154-300 cells per sample. Statistical analysis was done using Student’s T-test.

### Total lipid extraction

From 100 ml culture in each time point (ZT), we harvested 2ml aliquot and centrifuged it (4000 rpm, 10 min) in 20 ml glass vials. The supernatant was discarded and cell pellet equalized between the samples. The pellet was mixed with 1 mL of chloroform/methanol 2:1 by vigorous vortexing (2500 rpm, 30 min) and then centrifuged (4500 rpm, 10 min). 500 μL of resuspended pellet was transferred to clean 2 ml glass vial, vortexed (2500 rpm, 20 min) with 200 μL of LC-MS-grade water to extract polar molecules and left for gravity phase separation. The lower, organic layer containing lipids was transferred to clean 2 ml GC-MS glass vial and evaporated to dryness in a GeneVac evaporator, being finally ready for analysis by GC-MS.

### GC-MS

GC-MS analyses were performed on an Agilent 7890B GC system coupled to an Agilent 5977A Mass Selective Detector, using a Phenomenex Zebron ZB5-HT Inferno (35m × 250 μm × 0.1 μm). The TAGs and other fatty acid esters contained in the dry samples were resuspended and derivatized with methanol into fatty acid methyl esters via alkali catalysis in an automated fashion before the injection of each sample. The injector and interface were operated at 275 °C and 230 °C, respectively, and the oven heated from 50 °C to 260 °C at a rate of 7 °C min^-1^ and then isothermally held for 5 minutes. The total run time for each sample was 37 min. Helium was used as carrier gas, with a flow of 2 ml min^-1^. 1 μL of sample was injected in splitless mode. The MS was operated in scan mode (50-500 Da), with an ionization energy of 70 eV.

## Results

### Establishment of stress memory growth setup

As a first step, we sought to establish stress memory growth protocol for *C. reinhardtii*. We optimized light intensity conditions (Fig.S1a) and tested culture growth in medium depleted of nitrogen source or in default TAP medium with standard nitrogen source concentration (Ngan *et al.*, 2015). As expected, our test showed substantial doubling and increased OD600 for cultures grown in default medium, but not in the nitrogen-depleted samples (Fig.S1b).

Data published previously suggests that lipid accumulation in stressed *C. reinhardtii* cells reaches substantial levels already after 2 day-long growth under nitrogen depletion without lethal effect in the culture (Ngan *et al.*, 2015). Therefore, we designed stress priming setup, allocating 2 days for stress treatment and 2 days for the recovery period (Fig.1a). We set samples to be treated with two nitrogen depletion events (stress primed) or single stress event (control), harvesting aliquots for further analyses at timepoints (*zeitgeber*): ZT2 and ZT4, as indicated on the scheme (Fig.1a). We used 3 biological replicates for each timepoint in nitrogen-deprived and nitrogen-control samples and used harvested aliquots for microscopy or gas chromatography coupled with mass spectrometry (GC-MS/MS) (Fig.1b). To exclude the possibility that 1st nitrogen depletion event interferes with a readout from 2nd nitrogen depletion and ensure constant cell growth, we washed cultures with TAP-N (nitrogen depleted medium) before each stress event and split the cells 1:1 directly after 1^st^ stress (after ZT2) and after recovery window - directly before 2^nd^ stress (before ZT4). Our OD600 measurements suggested that the cells were actively dividing after nitrogen depletion events (Fig.1c).

**Fig.1.**
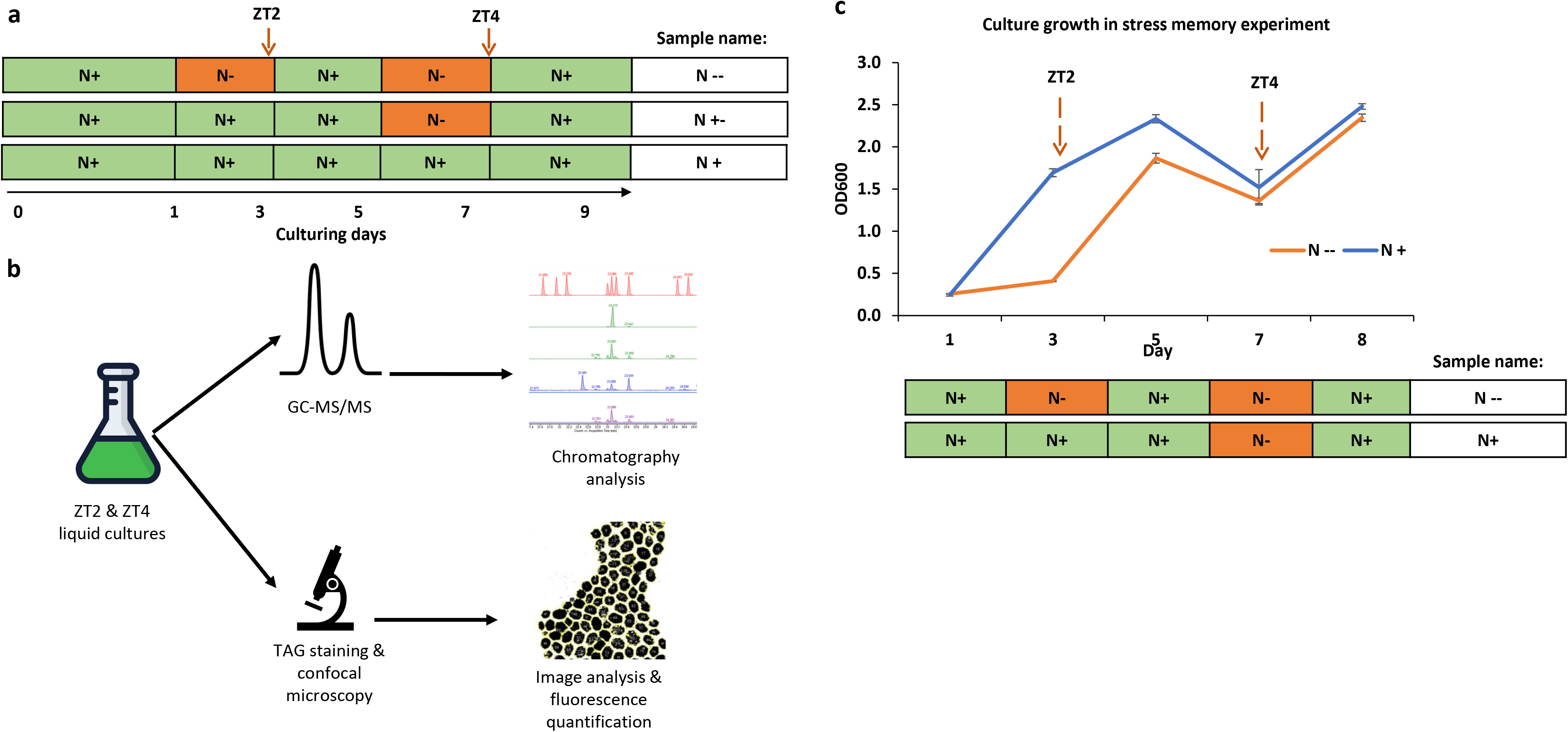
Stress memory growth schematic and cell growth. **a.** Stress memory growth scheme. ZT (*zeitgaber*) indicate harvesting timepoints for TAG measurements. N+ and N-labels correspond to nitrogen-control and nitrogen-depleted growth conditions., respectively. **b.** Post-harvesting workflow schematic **c.** Growth measurements (OD600) of cells in stress memory experiment. Error bars correspond to standard error from 3 biological replicates (separate flasks).

### Optimization of TAG staining and image analysis workflow

Using previously harvested aliquots at indicated timepoints, we sought to measure TAGs accumulation through histochemistry, microscopy and image analysis. Firstly, we performed chemical fixation of cells and their Nile Red staining of TAGs (see methods). Nile Red is an efficient TAG marker and its fluorescence was reported to be tightly correlated with TAG concentration in *C. reinhardtii* (Johnson *et al.*, 2017). Afterwards, we optimized confocal microscopy conditions (see methods) to separate Nile Red signal from cellular autofluorescence and acquired Z-stacks of prepared specimens to capture Nile Red signal in 3D. We observed only background autofluorescence in negative microscopy controls (mock-stained nitrogen-depleted (N-) or Nile Red-stained nitrogen-control (N+) samples) without specific Nile Red signal (Fig.2a). In contrast, Nile Red-stained nitrogen-depleted specimen showed TAGs’ accumulation in lipid bodies, as reported previously (Wang *et al.*, 2009), and selective Nile Red signal (Fig.2a), which confirmed specificity of staining and microscopy workflow.

**Fig.2.**
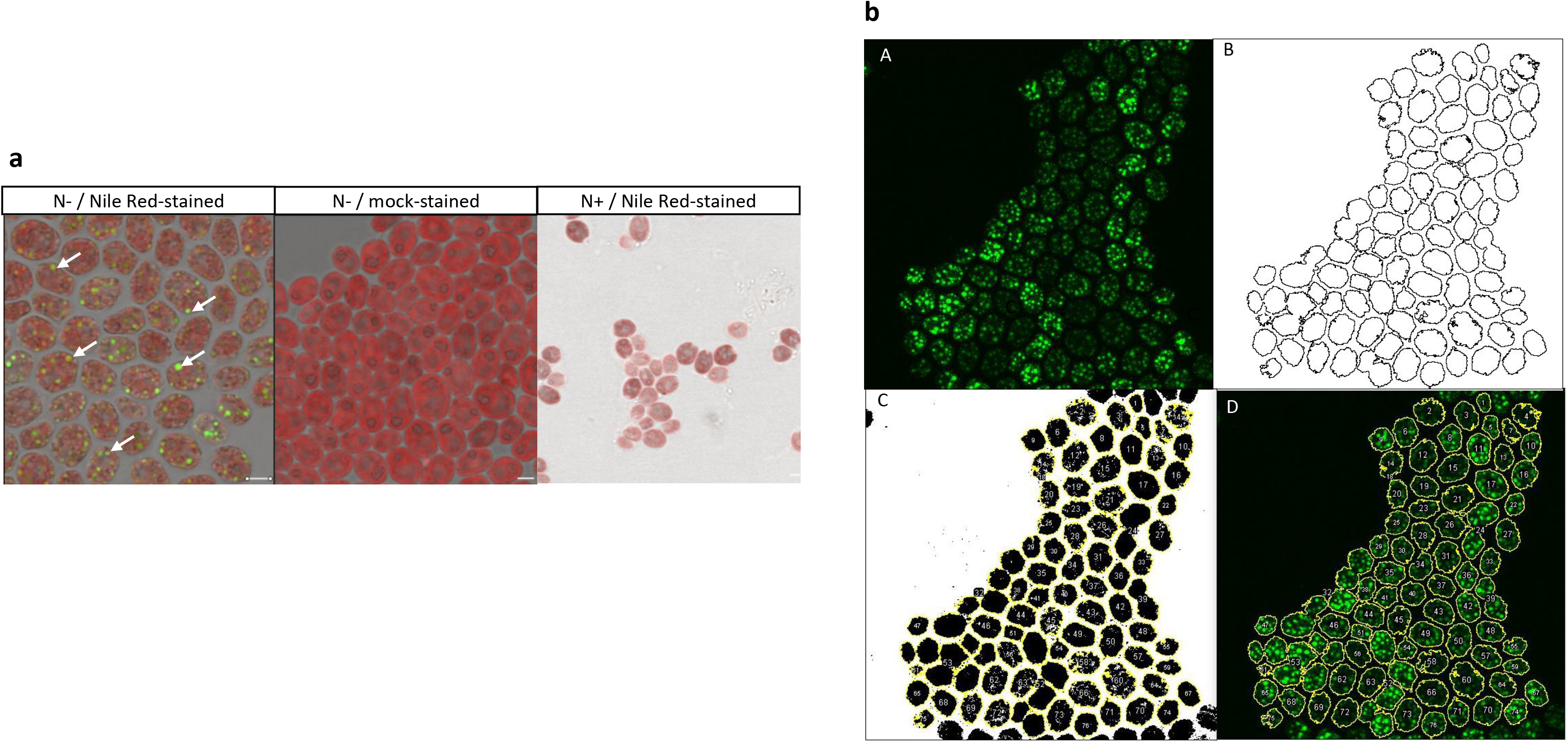
TAG staining and microscopy workflow. **a.** Exemplar images for Nile Red- and mock-stained samples. N-label corresponds to nitrogen-depleted samples, whereas N+ to control with standard nitrogen source concentration. The arrows correspond to examples of lipid bodies specifically stained by Nile Red. **b.** Image analysis workflow: A) Z-projection for Nile Red-specific channel; B) cell segmentation with pixel intensity thresholding; C) cell annotation on binary image D) cell annotation on original image for intensity quantification. Scale bar corresponds to 5 μm.

To properly quantify Nile Red-stained TAG accumulation, we established a workflow for image analysis. Firstly, we converted Z-stack to single images by slice summing in Z-projection. Afterwards, we segmented cells with minimum error threshold to create masks used as regions-of-interest (ROIs) outlining cell borders. Finally, we quantified mean fluorescence intensity in grey value inside ROIs as a way to measure Nile Red-stained TAG intensity. The examples of different workflow stages are present in Fig. 2b.

### TAG intensity quantification

Following measurement of a raw Nile Red-stained TAGs’ fluorescence intensity, we normalized technical differences between the images. To this end, we quantified background fluorescence outside the cells on the image and subtracted its value from the specific Nile Red signal from within the cells. We used normalized intensity measurements compare stress-primed sample at ZT4 (2^nd^ stress exposure) with its controls: stress-memory samples at ZT2 (1^st^ stress exposure, ZT2 N --) and single-stressed samples at ZT4 (ZT4 N +-). To ensure statistical power of the analysis, each sample had 3 biological replicates and the intensity measurements in each biological replicate was done on 154-300 cells.

As a result, we observed a clear and significant difference between stress-primed samples and single-stressed controls. Stress-primed samples showed downregulation of TAGs’ fluorescence intensity (Fig.3a, b), suggesting a differential response to recurrent nitrogen deprivation conditions and a memory effect of previously encountered stress. We do not envisage that an observed effect at ZT4 (2^nd^ stress) is a technical bleed-through from ZT2 (1^st^ stress), since cells has been split twice before ZT4 and the cultures were actively dividing throughout the experiment (Fig.1c). In turn, this observation in such experimental setup suggests a memory effect that is heritable over cell division and potentially of epigenetic nature.

**Fig.3.**
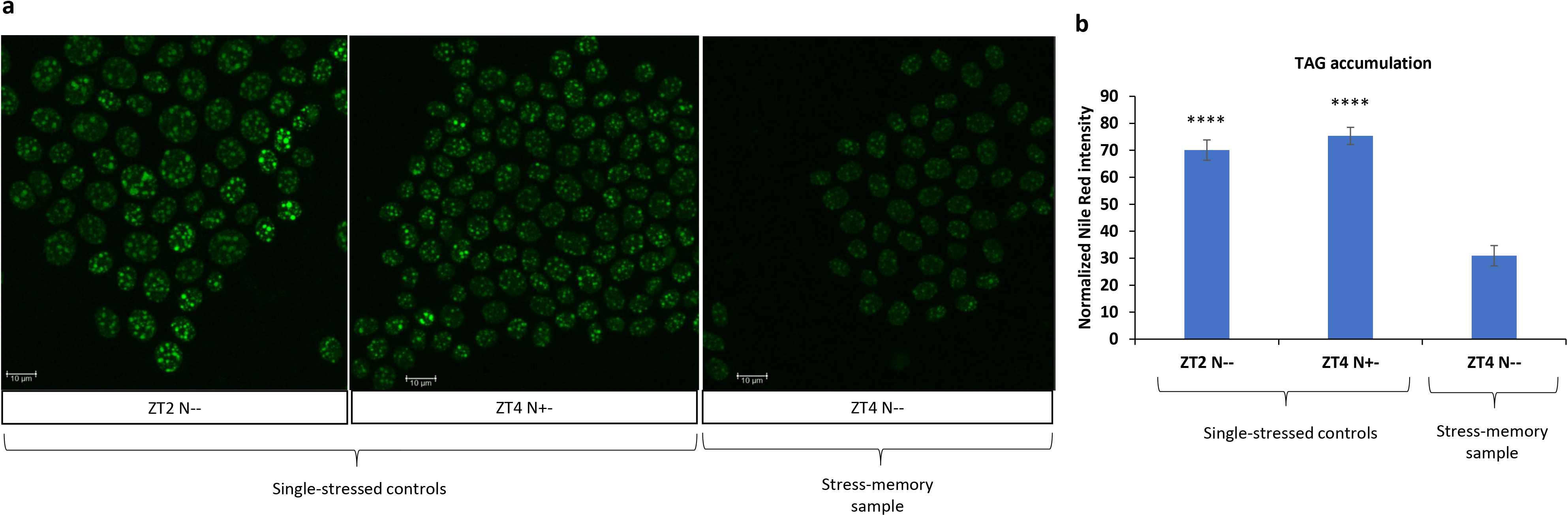
Imaging of TAG accumulation. **a.** Representative images of TAG-stained cells from stress-memory samples (N --) and single-stressed controls (N +- and N – at 1^st^ nitrogen depletion). Scale bar corresponds to 10 μm **b.** Nile Red intensity quantification as a proxy for TAG accumulation measurements. Error bars correspond to SEM from 3 biological replicates and statistics were calculated using Student’s t-test in comparison: ZT2 N– to ZTN– and ZT4 N+- to ZT4 N--.

### TAG profiling using GC-MS/MS

To qualitatively assess the lipid composition of the accumulated TAGs and complement the confocal microscopy measurements, we performed a gas chromatography mass spectrometry (GC-MS) lipidomic analysis. We extracted lipids from stress-primed cultures (ZT4 N-/-) and single-stressed control harvested at the same time (ZT4 N+/-). The extracted lipids underwent transesterification to generate volatile fatty acid methyl esters (see methods), which were subsequently analysed by GC-MS, along with standard fatty acid libraries.

Our results reveal that the qualitative profile of lipid types did not show substantial differences between stress-primed samples and single-stressed controls (Fig.4), despite different lipid accumulation observed through histochemistry. This observation suggests that quantitative stress priming response in TAGs accumulation does not come with qualitative change in lipid composition.

**Fig.3.**
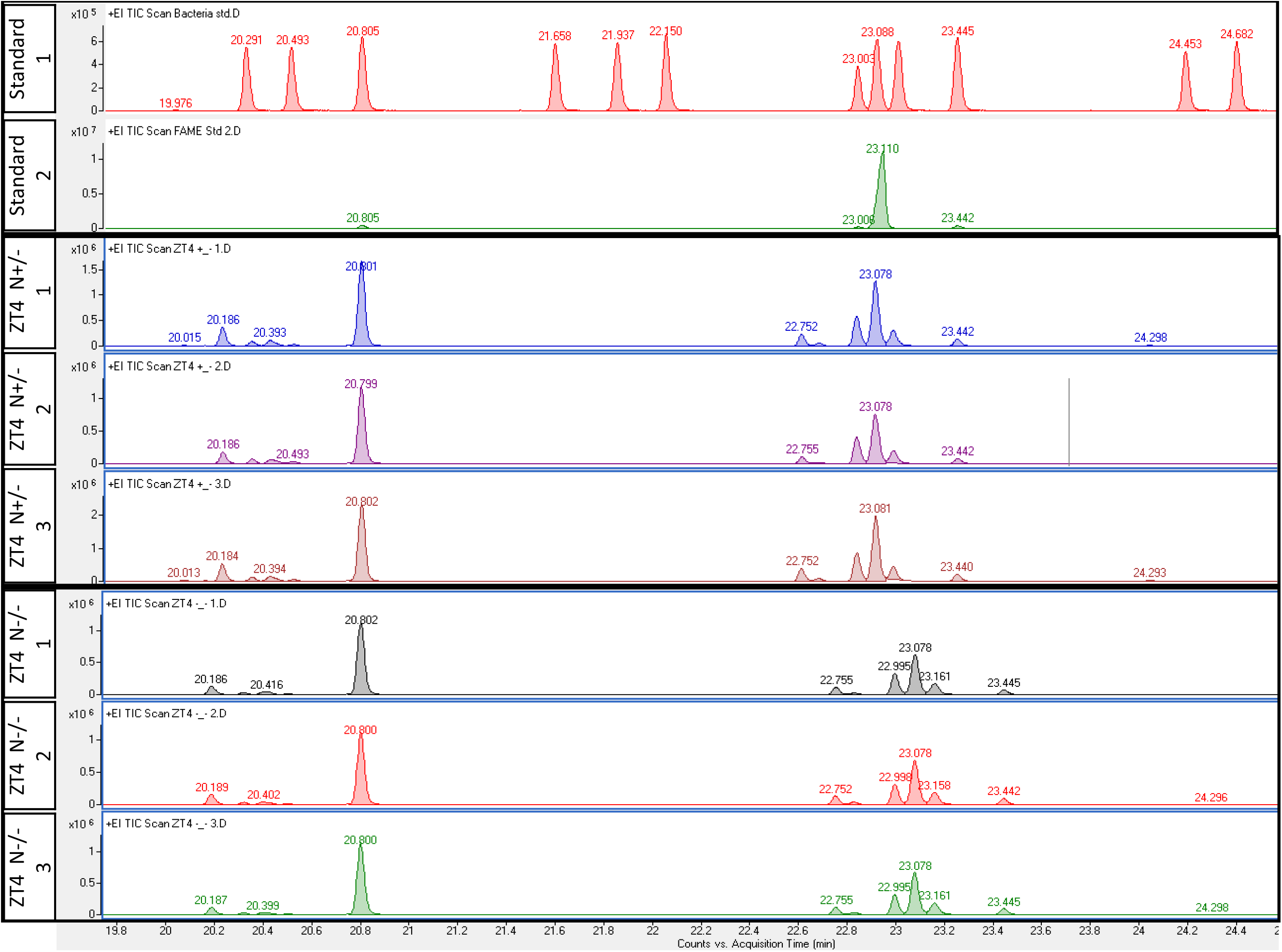
TAG profiling through GC-MS/MS. GC-MS TAG profiling. Chromatograms depict retention peaks of single-stressed ZT4 N+- and stress-primed ZT4 N– samples, each in 3 biological replicates as indicated in the numbers on the side labels. Two top bars correspond to GC-MS/MS lipid standards.

## Discussion

In summary, we observed a differential TAG accumulation caused by stress memory in Chlamydomonas cells. Cells subjected to two rounds of nitrogen depletion events show attenuated TAG accumulation, compared to single-stressed counterparts. Interestingly, this change was observed on quantitative level, as judged from histochemical measurements, but not in qualitative fashion – stress-memory samples do not exhibit substantial differences in the abundance of lipid subtypes, comparing to the controls. TAGs quantitative downregulation in stress-memory samples indicates that cells exhibit attenuated nitrogen deprivation response based on the previous stress exposure or respond to further stress events with pathways antagonistic to TAGs’ accumulation. Given the ongoing growth of the culture throughout the experiment, we envisage that observed response in stress-memory samples is potentially of epigenetic nature. Future steps should aim to elucidate the mechanism of stress memory and identify its causal factors.

In conclusion, our results highlight that metabolic stress memory of nitrogen depletion exists in model green alga and suggests that metabolic stress memory mechanisms could be widely conserved in non-vascular plants.

## Acknowledgements

We would like to thank Prof. Alison Smith (University of Cambridge, UK) and her group for providing

*C. reinhardtii* culture stocks and salt solutions for TAP medium; Prof. Jim Hasselof (University of Cambridge, UK) for organizing OpenPlant-Biomaker forum; Dr Andrew Truman (John Innes Centre, UK) for holding the grant and Paul Brett (John Innes Centre, UK) for the help with GC-MS/MS. This work was supported by Biomaker-OpenPlant initiative funded by BBSRC and EPSRC as part of the UK Synthetic Biology for Growth programme. The authors declare no conflict of interest.

## Author contribution

PM and JSA designed the research and performed culture harvesting. PM performed TAG staining, microscopy, image analysis and intensity quantification. JSA performed lipid extraction, gas chromatography and spectrometry peak analyses. PM, together with JSA, wrote the manuscript.

## Conflict of interest

The authors declare no conflict of interest.

## Notes

### Competing Interest Statement

The authors have declared no competing interest.

