## Supplementary figure 1 for "*Chlamydomonas reinhardtii* exhibits stress memory in the accumulation of triacylglycerols induced by nitrogen deprivation"

**a**

| Light setting | Measurement average |
|---|---|
| 5LS | 245 ± 11.9 |
| 4LS | 144 ± 8.29 |
| 3LS | 53 ± 2.94 |
| 2LS | 35 ± 5.35 |
| 1LS | 18 ± 2.83 |
| | unit: umol m-2 s-1 |

**b**

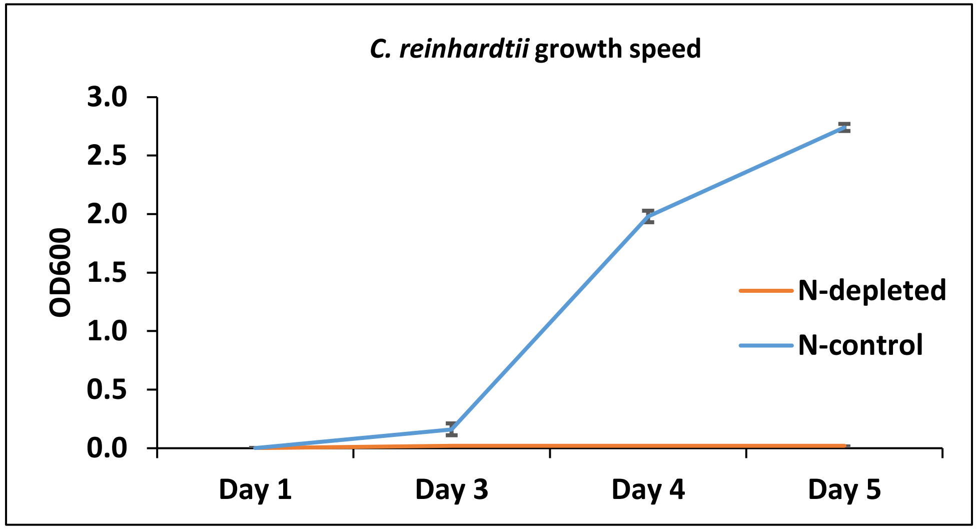


**Fig.S1 Growth conditions' optimization. a.** Light measurements in different settings of Sanyo 352-MLR-352-PE cabinets. Values indicated the average of 3 measurement in different positions on the growth chamber shelf. Plus-minus sign is followed by standard deviation from these measurements. Highlighted in red is the setting used for the further steps. **b.** Growth speed measurements in cells under nitrogen-control (in default TAP medium) or nitrogen-depleted conditions (TAP medium without nitrogen source). Error bars correspond to standard error from 2 biological replicates.
